# Multi-omics analysis of AML cells treated with azacitidine reveals highly variable cell surface proteome remodeling

**DOI:** 10.1101/369322

**Authors:** Kevin K Leung, Aaron Nguyen, Tao Shi, Lin Tang, Xiaochun Ni, Laure Escoubet, Kyle J MacBeth, Jorge DiMartino, James A Wells

## Abstract

Myelodysplastic syndromes (MDS) and acute myeloid leukemia (AML) are diseases of abnormal hematopoietic differentiation with aberrant epigenetic alterations. Azacitidine (AZA) is a DNA methyltransferase inhibitor (DNMTi) widely used to treat MDS and AML, yet the impact of AZA on the cell surface proteome has not been defined. To identify potential therapeutic targets for use in combination with AZA in AML patients, we investigated the effects of AZA treatment on four AML cell lines (KG1a, HL60, HNT34, and AML193), representing different stages of differentiation. The effect of AZA treatment on these cell lines was characterized at three levels: the DNA methylome (methylation array), the transcriptome (gene expression array), and the cell surface proteome (glycoprotein capture with SILAC labeling). Untreated AML cell lines showed substantial overlap in their methylomes, transcriptomes, and cell surface proteomes. AZA treatment globally reduced DNA methylation in all cell lines, but changes in the transcriptome and surface proteome were subtle and differed among the cell lines. Transcriptome analysis identified five commonly up-regulated coding genes upon AZA treatment in all four cell lines, TRPM4 being the only gene encoding a surface protein, and surface proteomics analysis found no commonly regulated proteins. Gene Set Enrichment Analysis (GSEA) of differentially-regulated RNA and surface proteins showed a decrease in metabolism pathways and an increase in immune defense response pathways. As such, AZA treatment in four AML cell lines had diverse effects at the individual gene and protein level, but converged to regulation of metabolism and immune response at the pathway level. Given the heterogeneous response of AZA in the four cell lines at the gene and protein level, we discuss potential therapeutic strategies for combinations with AZA.

## Introduction

Myelodysplastic syndromes (MDS) and acute myeloid leukemia (AML) are hematopoietic malignancies that are genetically and epigenetically diverse in nature. As myeloid lineage cells differentiate from their hematopoietic stem/progenitor cells, aberrant epigenetic changes can occur at any differentiation stage, driving cells into cancerous phenotypes (1). As such, AML is routinely classified according to hematopoietic lineages by cell morphology or by cytometry using sparse antibody surface markers (2, 3). Among many epigenetic changes that occur in MDS and AML, the most well-characterized change is the DNA methylation of cytosine bases in CpG islands (4). In fact, a hallmark of epigenetic changes in AML is the redistribution of methylated CpG dinucleotides with loss of methylation across intergenic regions, primarily transposable elements and repeats, and gain of aberrant methylation near the promoters of a number of genes, including well known tumor suppressors such as p16^INK4a^ (5). As such, it is believed that these diseases are more sensitive to hypomethylating agents such as DNA methyltransferase inhibitors (DMNTi) (6, 7). One such DMNTi, azacitidine (AZA), has been efficaciously used for over a decade to treat MDS and AML (8, 9). At high doses, AZA induces rapid DNA damage and is cytotoxic; at lower doses, AZA induces DNA hypomethylation by covalent trapping and degradation of DNA methyltransferases, leading to loss of methylation in newly synthesized DNA (10, 11). It was recently shown that AZA treatment of cervical (12, 13) and colorectal (14) cancer cells can turn on the interferon response genes through reactivation of endogenous retroviruses (ERVs). This phenomenon, termed viral mimicry, is thought to induce anti-tumor effects by activating and engaging the immune system.

Although AZA treatment has demonstrated clinical benefit in AML patients, additional therapeutic options are needed (8, 9). Our group has recently generated antibodies towards potential targets in RAS-driven cancers and there is significant interest in identifying surface protein targets for antibody-derived therapeutic strategies in combination with AZA for treatment of AML (15). Currently, there are numerous antibody-based therapeutics in development for AML patients, targeting about a dozen cell surface proteins, but it is not clear if AZA changes the expression levels of these proteins (16). Furthermore, over 15 ongoing clinical trials are investigating the combination of AZA and checkpoint inhibitors in various leukemias and solid tumors, since AZA induces checkpoint inhibitory molecules on both tumor and immune cells (7, 17). To identify cell surface markers, cell surface capture proteomics has recently emerged as a highly sensitive target discovery technology and has been used to define a large number of common and distinct markers in AML (18, 19). Taken together, a broader understanding of how AZA treatment remodels the cell surface proteome in AML cells could aid in identifying surface protein targets for antibody-derived immune therapy, leading to unique immune-based therapies for use in combination with AZA.

Using a multi-omics approach, we characterized four AML cell lines, representing different stages of differentiation, and studied the changes in DNA methylation, RNA expression, and surface proteome induced by AZA treatment. Across the four cell lines, AZA reduced DNA methylation in nearly all of the hypermethylated CpG sites probed, but surprisingly the changes in gene expression and surface protein expression were few and diverse. Transcriptome analysis identified only one gene encoding a surface protein to be commonly up-regulated in all four cell lines, and surface proteomics analysis did not identify any commonly regulated proteins. Despite little overlap, functional analysis revealed some common responses among the four cell lines – down-regulation of genes and proteins in metabolism and up-regulation of genes in immune response. Collectively, our study detailed the distinct impact of AZA treatment in four AML cell line at the individual gene level, and illustrated that functional networks are commonly regulated.

## Results

### Methylome in AML cells and its regulation by azacitidine

Four well-characterized AML cell lines, KG1a, HL60, HNT34, and AML193, were chosen to reflect a gradient of differentiation stages along the myeloid lineage, according to the French-American-British (FAB) classification system). KG1a (FAB M1) was established as a subline of KG1 myeloblasts with minimal maturation, but unable to differentiate (20, 21); HL60 (FAB M2) was established from a promyelocyte that can differentiate into neutrophils, granulocyte-like cells, or monocyte/macrophage-like cells by various chemicals (22); HNT34 (FAB M4) was established from chronic myelomonocytic leukemia (CMMoL) (23); and AML193 (FAB M5) was established from acute monocytic leukemia (24). The four cell lines also exhibited varied mutation profiles. *PHF6* (PHD Finger Protein 6) was the only gene mutated among all four cell lines, while genes like *TET1* (Tet Methylcytosine Dioxygenase 1), *DNMT3B* (DNA Methyltransferase 3B)*, and NRAS* (NRAS Proto-Oncogene) showed distinct mutation patterns in the four cell lines (**Supplemental Table 1**).

We determined the DNA methylome of each cell line using the Illumina Infinium EPIC array. The baseline DNA methylation profile for each cell line exhibited a bimodal distribution, representing hypermethylated and hypomethylated CpG sites (**Figure 1A**). The number of hypermethylated sites (beta value > 0.8) was highest for AML193 and lowest for HL60, following a general trend of increasing hypermethylation from the least to most differentiated cell line (**Figure 1B**). KG1a does not follow this trend of increasing baseline hypermethylation, potentially due to its differentiation from KG1 parental cell line and a deviation from the annotated early progenitor lineage. Comparing the methylation status across the four cell lines, 55% (342320/628240) of loci were commonly hypermethylated (beta value > 0.8) and 39% (108968/278579) of loci were commonly hypomethylated (beta value < 0.2) (**Figure 1B**). This indicates a high degree of similarity in the methylomes among the four cell lines.

**Figure 1.**
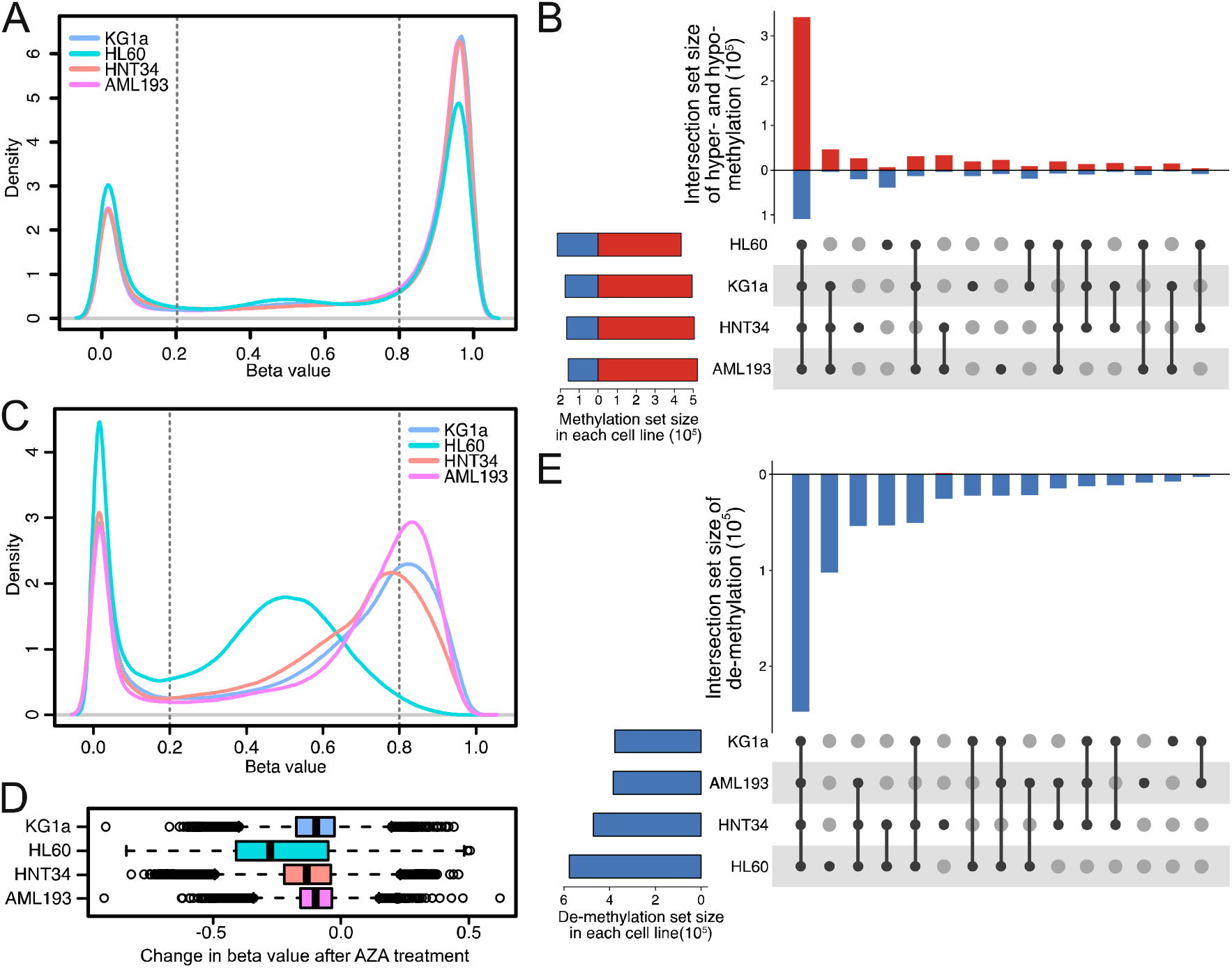
Azacitidine treatment drives global DNA de-methylation among all four AML cell lines. (A) Distribution with kernel density estimation for genome-wide beta values of the four cell lines exhibits a typical bimodal distribution in vehicle treated samples reflecting hyper- and hypomethylated genes. (B) Upset plot showing a high degree of overlap in all hypermethylated and hypomethylated sites among the four cell lines. In the top bar graph, overlapping hypermethylated sites (red) are indicated by the upward bars, and overlapping hypomethylated sites (blue) are indicated by the downward bars. The specific overlapping groups are indicated by the black solid points below the bar graph. Total hypermethylated and hypomethylated sites found in each cell line are indicated in the left bar graph. Hypermethylated sites were defined as probes with beta values > 0.8, and hypomethylated sites as beta values < 0.2. (C) Distribution with kernel density estimation for genome-wide beta values of the four cell lines in AZA treated samples shows drastic shifts of the hypermethylated peaks reflective of DNA de-methylation. (D) Box and whisker plot distribution of change in beta value after AZA treatment. Median change in beta value for KG1a (−0.097), HL60 (−0.28), HNT34 (−0.132), and AML193 (−0.099) shows AZA induced de-methylation. Box blot is defined by the first and third interquartile range, and whiskers extends up to 1.5 interquartile range above or below the box. (E) Global de-methylation across all cell lines shows a high proportion of common de-methylated sites (blue). Number of overlapping de-methylation sites are indicated by downward bars in the top graph and the specific overlapping groups are indicated by the black solid points below the bar graph. Total de-methylated sites found in each cell line are indicated in the left bar graph, showing that KG1a has the smallest de-methylation effect while HL60 has the largest effect reflecting differences in sensitivity to AZA treatment. De-methylation is defined as decrease in beta value > 0.1, and FDR adjusted p ≤ 0.05.

To study the effects of AZA treatment on the regulation of DNA methylation, each of the four cell lines was treated with AZA (0.5 μM) for 3 days, followed by a four-day drug holiday (**Supplemental Figure 1A).** With this treatment regimen, AZA inhibited cell growth to varying degrees in the AML cell lines, with ~85% growth inhibition in AML193 cells (most sensitive) and ~30% inhibition in HL60 cells (least sensitive) **(Supplemental Table 2)**. Remarkably, AZA treatment reduced methylation in nearly all of the hypermethylated sites probed (**Figure 1C-D, Supplemental Figure 2**). The median change in methylation cross all CpG sites ranged from -0.097 for KG1a cells to -0.28 for HL60 cells (**Figure 1D)**. The greater reduction of DNA methylation seen in HL60 compared to the other three cell lines could be due to a lower basal expression of the *de novo* DNA methyltransferases (DNMT3A and DNMT3B) in the HL60 cells, as detected in the transcriptome data **(Supplemental Figure 3**). Comparing the effects of AZA across all four cell lines, a large proportion of CpG sites with reduced DNA methylation were shared among the four cell lines (37% or 247,715 shared out of 657,868 total) (**Figure 1E**). The loci with reduced methylation induced by AZA were uniformly distributed across the genome **(Supplemental Figure 4).** Further Gene Set Enrichment Analysis (GSEA) of the AZA-regulated DNA methylation loci did not identify any functionally enriched gene sets (data not shown).

### Transcriptome in AML cell lines and its regulation by azacitidine

As DNA methylation at promoter CpG islands can be associated with transcriptional silencing, we next assessed RNA expression profiles of the four cell lines in the absence and presence of AZA treatment. Using a similar approach as for comparing baseline DNA methylation status, highly- and lowly-expressed genes were defined by the expression level from the highest and lowest tertiles in each cell line. At baseline, 53% (10971 out of 20517 total) of all high- and low-expressing genes were common among all four cell lines (**Figure 2A**). Despite a shared gene expression profile, functional analysis using Gene Set Variation Analysis (GSVA) with a hallmark set of key pathways indicated that each AML cell line had a distinct biological state. Specifically, cell-cycle related genes were highly expressed in AML193 and KG1a cells, MYC pathway genes in KG1a and HL60 cells, and TGFβ, WNT, and Notch signalling genes in HNT34 cells (**Figure 2B**). Similarly, GSVA using an expanded gene set list of GO term analysis also revealed differences in differentiation and cell death activation pathways (**Supplemental Figure 5**).

**Figure 2.**
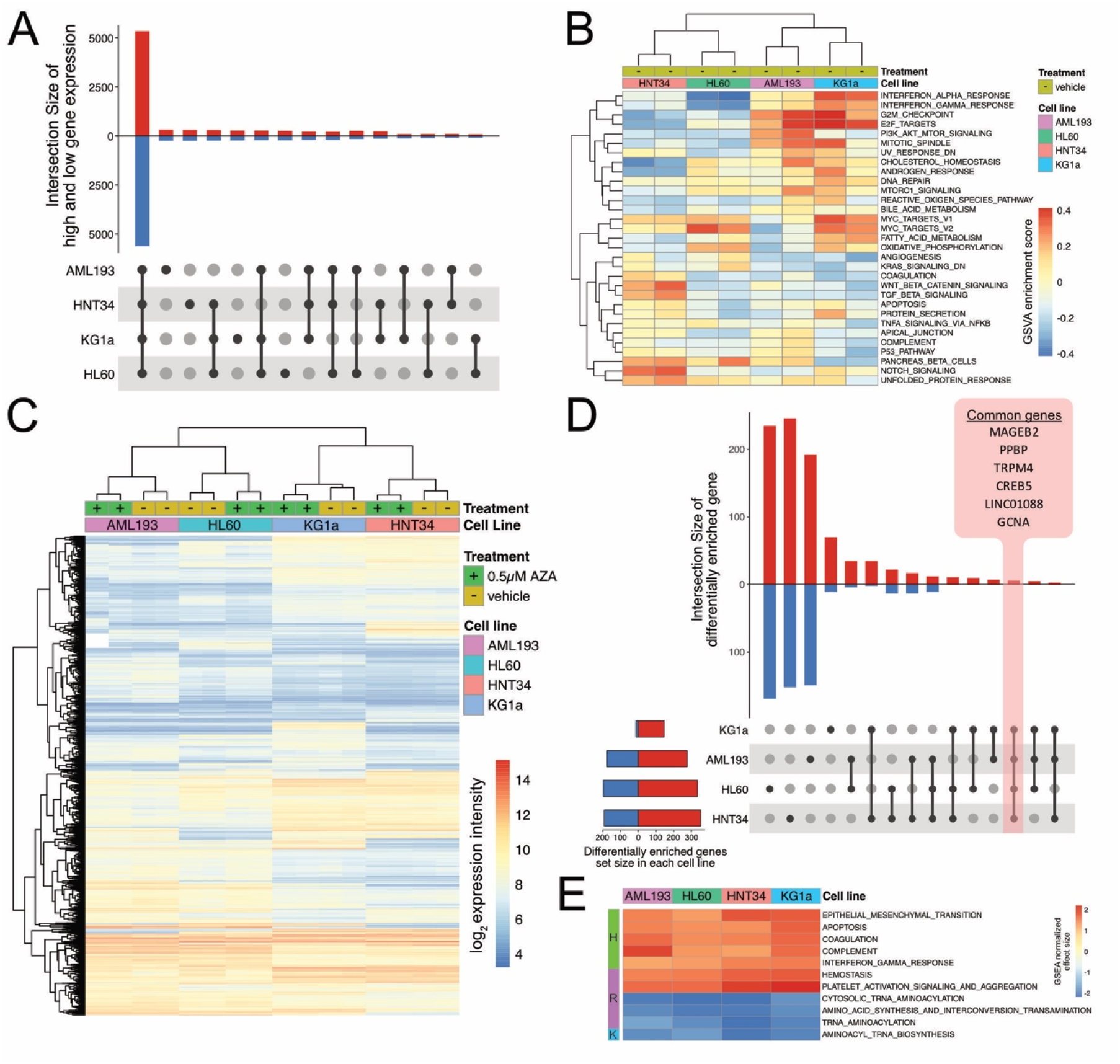
Azacitidine treatment induces unique and subtle transcriptome changes in the four AML cell lines. (A) Overlap of the distribution of high and low RNA expression at baseline levels. In the top bar graph, overlapping highly expressed genes (red) are indicated by the upward bars, and overlapping lowly expressed genes (blue) are indicated by the downward bars. The specific overlapping groups are indicated by the black solid points below the bar graph. High expression is defined as log_2_ expression intensity in the top tertile, while low expression is defined as log_2_ expression intensity in the bottom tertile (n = 20517 unique genes mapped from > 54000 probesets). (B) Gene Set Variation Analysis of the four cell lines at baseline levels (variation score > 0.2 in at least one cell line). GSVA was performed using the MSigDB hallmark gene set. (C) Hierarchal clustering of top variable gene expression profiles (IQR of probesets between four cell lines > 1, n = 5747). The differences in gene expression varied more among cell lines than by AZA treatment. (D) Most of differentially regulated genes induced by AZA are unique to each cell line, and only six genes were commonly up-regulated by AZA among the four cell lines. In the top bar graph, up-regulated genes (red) are indicated by the upward bars, and down-regulated genes (blue) are indicated by the downward bars. The specific overlapping groups are indicated by the black solid points below the bar graph. Total differentially regulated genes for each cell line are indicated in the left bar graph showing that transcriptome of KG1a was least affected by AZA treatment, while transcriptome of HNT34 was most affected. Up-regulated (red) and down-regulated (blue) genes are defined as genes with fold change > 2 and adjusted p value < 0.05. (E) Biological processes are commonly enriched in all four cell lines upon AZA treatment despite small overlap in specific genes. Gene Set Enrichment Analysis (GSEA) was performed using Hallmark (H), Reactome (R), and KEGG (K) pathways. Significant pathway was defined as GSEA normalized effect size > 1 or < -1. An extended GSEA of significantly enriched gene sets in at least one cell line is presented in supplemental Figure 6A-B.

Upon AZA treatment, only a small percentage (~0.8% to 2.4% of all genes) of the transcriptome was regulated in each AML cell line, despite the large changes observed in the DNA methylome. Hierarchical clustering of the top variable gene expression profiles showed clustering by cell line and not by AZA treatment, suggesting that AZA did not have a dominant effect on the transcriptome (**Figure 2C)**. The number of differentially regulated genes varied from ~160 genes for KG1a cells to ~500 genes for HL60 and HNT34 cells (**Figure 2D**). The changes in gene expression per cell line were much lower compared with those observed for DNA methylation (45 to 70% of all CpG sites were significantly demethylated). The effects on gene expression were also much more stochastic and appeared to be divergent among the cell lines, with only five coding and one non-coding gene uniformly up-regulated in all four cell lines (**Figure 2D**). The commonly up-regulated coding genes were *TRPM4* (Transient Receptor Potential Cation Channel subfamily M member 4), *PPBP* (Pro-Platelet Basic Protein), *MAGEB2* (MAGE family member B2), *CREB5* (cAMP Responsive Element Binding Protein 5), and *GCNA* (Germ Cell Nuclear Acidic Peptidase). The non-coding gene *LINC01088* found to be up-regulated in all four cell lines was recently implicated as a tumor suppressor found in reduced levels in ovarian tumors and acts by targeting miR-24-1-5p mediated regulation PAK4 expression(25) . Despite the unique gene regulation in each cell line, functional level analysis with GSEA showed several common pathways regulated by AZA in all four cell lines (**Figure 2E, Supplemental Figure 6A-B**). The enriched gene sets included increased expression of epithelial mesenchymal transition, apoptosis, coagulation, complement, interferon gamma response, hemostasis, platelet activation signaling and aggregation; and decreased expression of tRNA aminoacylation and amino acid synthesis. In general, these gene sets represent activation of immune response and repression of metabolism.

### Unique cell surface proteome regulation by azacitidine

In an effort to identify novel therapeutic targets induced by AZA, we probed the surface proteomes of the four cell lines using a modified cell-surface-capture of N-linked glycosylated proteins protocol and quantified the changes induced by AZA treatment using stable isotope labeling by amino acids in cell culture (SILAC) (26, 27) **(Supplemental Figure 1B)**. Each experiment was performed in both the forward and reverse SILAC mode, and pairwise comparisons of the enrichment ratios for the biological replicates showed good reproducibility (**Supplemental Figure 7A-B**). To compare the baseline surface protein profiles among the four cell lines, we extrapolated a subset of proteomics data from heavy-labeled vehicle-treated cells (**Supplemental Table 3**). At baseline, a total of 875 unique surface membrane proteins were identified among the four cell lines, and a common set of 232 proteins were detected in all four cell lines (**Figure 3A**). Historically, classification of cell lineage is dependent on staining of CD markers on cells. Here, we detected an extensive number of CD markers expressed on the four cell lines and reported an estimate of protein abundance (**Figure 3B).** Among the common CD markers identified were several therapeutic targets of AML, such as SIGLEC-3 (CD33) and integrin associated protein (CD47) (16). Comparison between our unbiased proteomics and transcriptome data to known immunophenotyping CD markers of the four cell lines also showed remarkable agreement (**Supplemental Table 4**).

**Figure 3.**
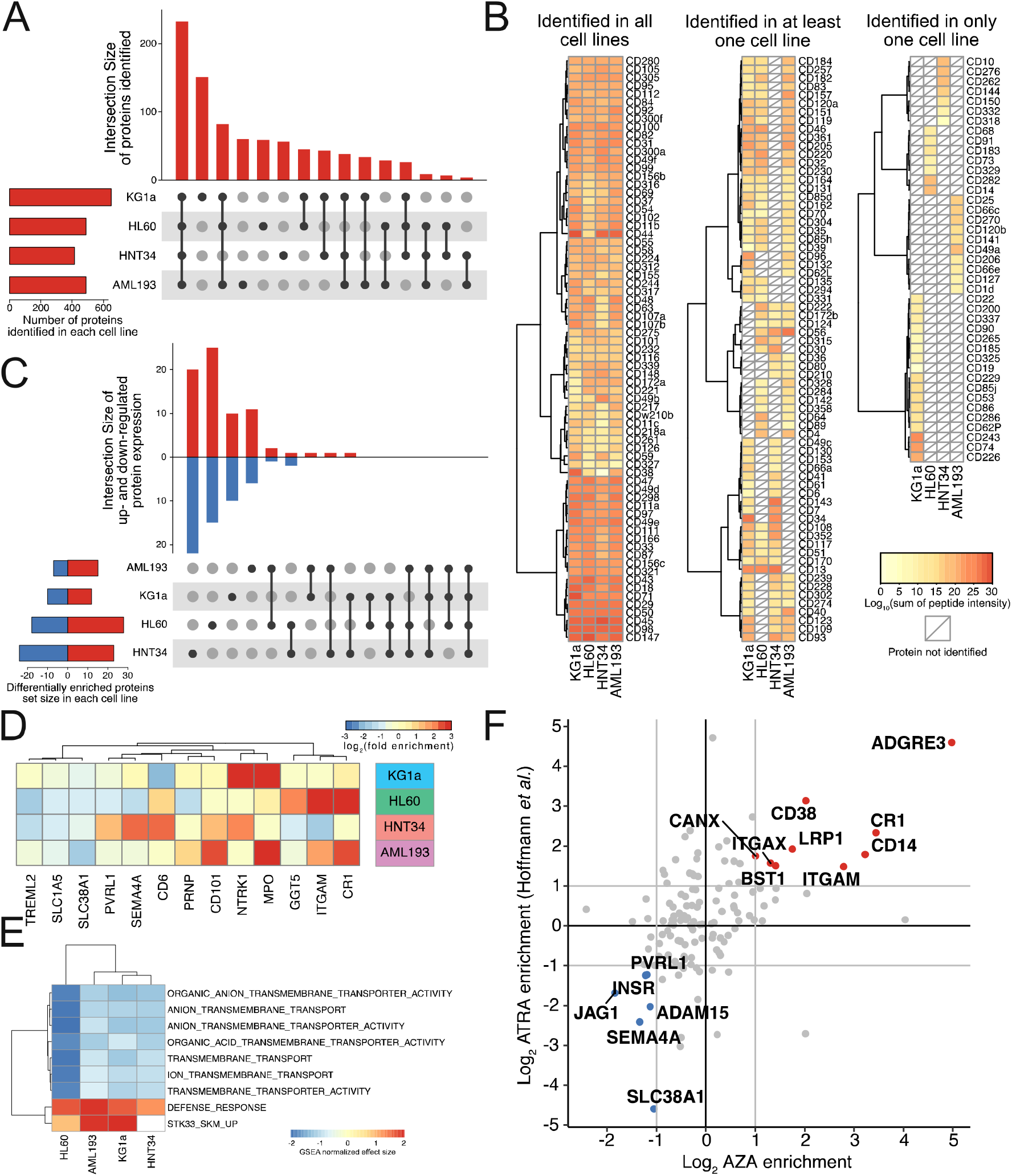
Surface proteome changes induced by azacitidine treatment in the four AML cell lines. (A) Upset plot showing an overlap of 232 commonly identified surface proteins in the four AML cell lines in vehicle treated samples. Number of overlapping protein identification are indicated in the top bar graph and the specific overlapping groups are indicated by the black solid points below the bar graph. Total surface proteins identified in each cell line are indicated in the bottom left bar graph. Protein identification was defined by detection of at least one peptide belonging to vehicle treated cells labeled with heavy isotope, filtered by having at least one well-quantified peptide defined by Skyline’s isotope dot product of > 0.8. (B) CD markers identified by cell surface capture enrichment in vehicle treated sample. Heat map is separated by CD markers identified in all four cell lines (left), in at least one cell line (middle), and only in one cell line (right) and shaded from yellow to red to reflect estimated abundance. Estimation of abundance was based on the logarithmic sum of well-quantified peptide intensities for each protein. (C) AZA induced unique changes on the surface proteome. In the top bar graph, overlapping up-regulated proteins are indicated by upward bars, and overlapping down-regulated proteins are indicated by downward bars. The specific overlapping groups are indicated by the black solid points below the bar graph. Total differentially regulated surface proteins for each cell line are indicated in the left bar graph showing that the surface proteome of KG1a and AML193 were least affected by AZA treatment, while HL60 and HNT34 were most affected. No commonly regulated protein was identified among the four cell lines from surface proteomics data. Up-regulation was defined as median SILAC ratio > 2, p value < 0.05, and downregulation was defined as median SILAC ratio < -2 in blue. (D) Proteins with significant changes in at least two cell lines are shown to illustrate that no single surface proteins is commonly regulated by the treatment of AZA (n=13). (E) Top enriched gene sets affected by AZA treatments identified by Gene Set Enrichment Analysis (GSEA) of the proteomics dataset using GO term analysis shows there were common functional changes despite few specific proteins that overlapped. (F) Comparison of surface proteomics data between AZA treatment vs ATRA treatment on HL60 cells. Pearson correlation between the two datasets was 0.44. Data for ATRA treatment in HL60 was obtained from Hofmann *et al.* (18).

We next explored how AZA treatment affected the surface expression of membrane proteins using SILAC quantification (**Supplemental Figure 1A**). The number of differentially regulated proteins ranged from 22 to 47, or 5 to 10% across the cell lines, and no protein was commonly regulated upon AZA treatment (**Figure 3C**). In fact, the majority of the changes were unique to each cell line, and only 13 proteins were significantly differentially regulated in at least two cell lines **(Figure 3D)**. Some proteins, such as CR1 (Complement C3b/C4b Receptor 1), ITGAM (Integrin Subunit Alpha M), and MPO (Myeloperoxidase) appeared to be generally up-regulated by AZA treatment, but was statistical significance in only 2 or 3 cell lines. Western blot validation of the regulation of ITGAM (CD11b), a common marker of neutrophil/monocyte differentiation, was consistent with SILAC quantification (**Supplemental Figure 8**). Specifically, expression of ITGAM was up-regulated in HL60 and AML193 cells, did not change in KG1a cells, and was down-regulated in HNT34 cells. Together, the surface proteomics analysis, similar to gene expression analysis, suggests that the effects of AZA are largely dependent on the inherent differences among the AML cell lines. Hierarchical clustering of significantly enriched protein expression showed a distinctive protein regulation profile in each cell line (**Supplemental Figure 7C**). Further functional analysis using GSEA with GO terms indicated an increase in immune response, and a decrease of various transmembrane transporters (**Figure 3E**). The pathway analysis of proteomics was consistent with the pathway analysis of RNA, showing activation of immune response and repression of metabolism.

Previously, Hofmann *et al*. used three cell surface capture techniques (CSC) to identify ~500 total surface proteins between two AML cell lines (HL60 and NB4) that represent the M2 and M3 stages of AML (18). Indeed, comparison of our HL60 datasets showed an overlap of 230 identified proteins (**Supplemental Figure 9**). To understand cellular differentiation, Hofmann *et al.* further characterized the surface proteome in response to all-trans retinoic acid (ATRA). Despite different mechanism of action, both ATRA and AZA treatment of HL60 cells are known to induce granulocytic and monocytic differentiation. To this end, we compared our HL60 dataset to the existing dataset and observed a considerable overlap of changes in the surface proteome (Pearson Correlation of 0.44, **Figure 3F).** Among the up-regulated proteins identified in both datasets are several known monocytic differentiation markers such as ITGAM (CD11b), CD14, and CD38, as well as some previously undefined markers such as ADGRE3 and CR1. It is remarkable that even though the two molecules target different cellular functions, a number of common targets emerged. As such, these proteins are potential therapeutic targets for subtypes of AML that undergo differentiation upon AZA or ATRA treatment.

### Comparisons of methylome, transcriptome, and surface proteome profiles

Having all three omics datasets allowed for comparisons of cellular states of the four cell lines with and without AZA at the DNA, RNA, and surface protein levels. Hierarchical clustering of each omics dataset primarily showed bigger variation among the individual cell lines compared to the variation due to AZA treatment (**Figure 4A**). At the DNA methylation level, KG1a (M1) clustered with HL60 (M2), while AML193 (M5) clustered with HNT34 (M4), consistent with their FAB classifications. At the transcriptome and surface proteome levels, however, KG1a (M1) and HNT34 (M4) clustered together, while AML193 (M5) and HL60 (M2) clustered together. KG1a and HNT34 are cell lines known to be non-responsive to differentiation agents such as GM-CSF (20, 23), while AML193 and HL60 have been shown to differentiate in response to various reagents (22, 28). Therefore, the four cell lines defined by cellular morphology may correlate with the methylome, but the functional state of these cells are more closely correlated to transcriptome and cell surface proteome.

**Figure 4.**
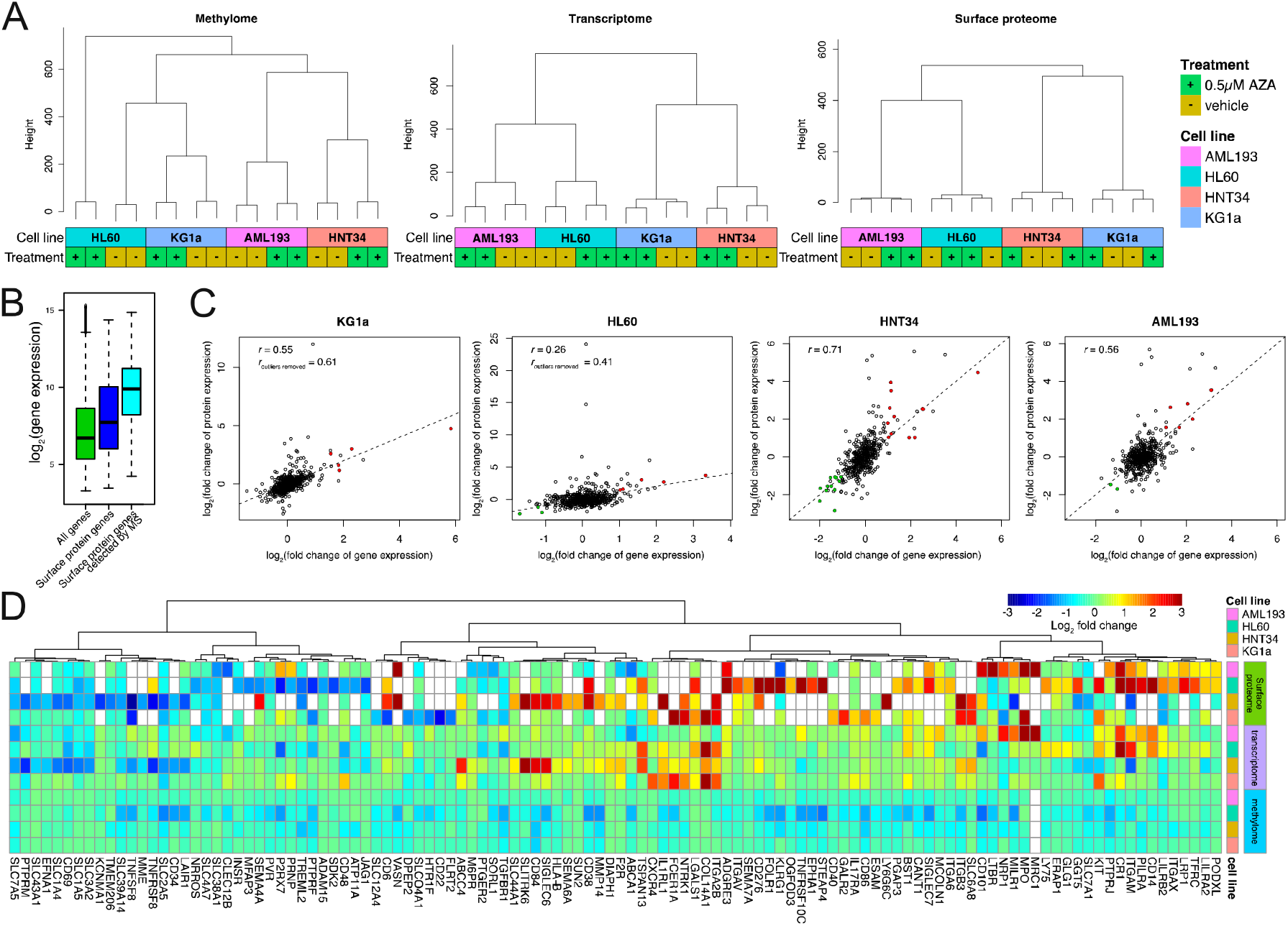
Omics comparison between methylome, transcriptome, and surface proteome in AML cells treated by AZA. (A) Dendrogram of hierarchal clustering of methylome, transcriptome, and surface proteome data. In all cases, clustering was dominated by differences between cell lines rather than differences between vehicle and AZA treatment of any cell line. All experiments were performed in biological duplicate. (B) Representative RNA expression profile of all genes (green), of all genes annotated to be surface proteins (blue), and of genes identified by mass spectrometry experiment (cyan) illustrate that surface proteins have higher gene expression levels. RNA expression profiles for all cell lines are shown in **Supplemental Figure 9A**. (C) Correlation of changes in gene and protein expression. Pearson correlation (*r*) range from 0.2 for HL60 cells to 0.71 for HNT34 cells. Significantly up- and down-regulated genes and proteins are highlighted in red and green, respectively (p value < 0.05 for both gene and protein expression profile). Dashed lines (y = x) are drawn for reference. (D) Changes in protein, gene expression, and DNA methylation for those genes with significant protein changes in at least one cell line are shown as a heatmap. Log_2_ fold changes were plotted for protein and gene expression, and average changes in beta values for CpG sites within 1500bp of transcriptional start site (TSS1500) of a given gene (also scaled to match the log2 FC scale of protein and gene expression changes). Proteins not identified are indicated in white.

Comparison between gene and surface protein expression showed that surface proteins identified in the proteomic datasets tend to have a higher transcriptional signal overall **(Figure 4B, Supplemental Figure 10A).** This likely reflects the nature of mass spectrometry in detecting proteins with the highest abundance. Quantitative comparison between the changes in RNA and protein expression after AZA treatment showed Pearson Correlation coefficients (*r*) ranging from 0.26 to 0.71 (**Figure 4C)**. Such correspondence has been previously reported in other whole cell proteomics experiments using cell lines (29-31). The differences between mRNA and protein levels are likely due to significant differences in regulation and steady-state turn-over.

Next, we compared the magnitude of changes in DNA methylation, gene expression, and surface protein expression in a subset of cell surface proteins that were significantly regulated in at least one of the cell lines (**Figure 4D, Supplemental Figure 10B**). Comparing the three omics datasets for these genes/proteins of interest, we found that while protein and transcript levels tracked reasonably well, changes in methylation status were not correlated with either transcript or protein level. Studies have shown that methylation effects on transcription can be both positive and negative (32, 33). This may explain why only a small subset of RNA changes can be directly accounted for by DNA methylation changes at either the corresponding gene promoter regions or gene body.

In an effort to identify AZA induced functional modules that might be missing from analyzing one type of data, we adopted a recently developed algorithm, SMITE (Significance-based Modules Integrating the Transcriptome and Epigenome) to further probe for the functional consequence that might be exerted jointly at methylation and transcription levels (34). Through combining the significance of gene regulation at both methylation and transcription levels and applying the joint scores onto a gene interaction network, this algorithm increases analysis power in identifying subnetworks of dysregulated genes with collateral information. Through this analysis, each AML cell line had 16-27 functional modules identified and these functional modules encompass 481 ~ 612 genes (**Supplemental Table 5**). In particular, analysis of the KG1a cell line benefited most from SMITE, where 528 genes were found to be dysregulated compared to 160 genes when analyzed using transcription data alone. This indicated wider functional impact of AZA treatment on the KG1a cell line when examining both sets of information in the context of gene interaction networks compared to transcriptomics alone. Also, when looking across all four cell lines, instead of the 5 common genes found to be transcriptionally dysregulated, SMITE identified 19 functional modules encompassing 310 genes. We observed recurrent themes of dysregulation of metabolism from multiple top functional modules identified across all four cell lines -HDC:HNMT, CTSA:GNS:IDS:NEU1, and PTGS2:LTC4S, representing genes involved in histamine metabolism, protein processing, and leukotrienes metabolism. Other top common functional modules affected were ‘natural killer cell mediated cytotoxicity’ and ‘osteoclast differentiation’. These findings were consistent with the impact of AZA on immune pathways and AML cell differentiation, and also identified novel functional modules. For example, HNMT encodes a methyltransferase that metabolizes histamine via N(tau)-methylation. Core genes from these functional modules would also serve as interesting therapeutic targets for follow up validation. AZA impacted different functional modules in the individual cell lines – top modules involved in various signaling pathways (mTOR, insulin, p53 etc) were affected in AML193 cells, while modules related to the MAPK signaling pathway was regulated in HL60 cells. The analysis here indicated that although methylation and transcription regulation by AZA might not be synchronized at the gene level across cell lines, functional networks appeared to be commonly regulated.

## Discussion

As omics technology becomes more widely accessible, integrating data analysis from orthogonal omics sources will be crucial to understand any biological question. In this study, we asked how AZA affects four different AML cell lines at three omics levels: the DNA methylome, RNA transcriptome, and surface proteome. This allowed us to compare the cancer cell lines at the epigenetic level and how that manifests into gene expression and surface protein expression. Our multi-omics study of the four AML cell lines showed that ~80% CpG sites, ~53% transcripts, and ~ 50% surface proteins overlap in methylation or expression pattern in vehicle treated cells. AZA treatment led to global reduction in DNA methylation, ranging from 45% to 70% of all probed CpG sites, while changes in mRNA and surface protein expression were much more subdued, ranging from 5% to 10%. One gene encoding a surface protein, *TRPM4*, was found to be commonly up-regulated by AZA treatment in all four cell lines, and may represent a potential novel therapeutic target for AML in combination with AZA. Comparing to previously published data of ATRA treatment in HL60, we identified several previously undefined markers such as ADGRE3 and CR1 that are potential therapeutic targets for subtypes of AML that undergo differentiation with AZA or ATRA treatment.

Despite relatively few changes observed at the transcriptome and surface proteome levels due to AZA treatment, functional analysis of RNA and protein regulation showed a general repression of metabolism and activation of immune response across the four cell lines. The repression of metabolism was consistent with the reduction of cell viability when treated with AZA, indicating some cytotoxic effects of AZA treatment (**Supplemental Table 2**). We also observe activation of immune responsive genes, which is consistent with previous studies showing that AZA treatment in cells of epithelial origin led to the transcription of endogenous retrovirus, and an induction of a number of immune response genes (AIM genes) related to anti-viral response(13). Even though most of the defined AZA-induced immune genes (AIM genes) were activated in the current study, the magnitude of induction was variable among all four cell lines (**Supplemental Figure 11**). Recently, a number of clinical trials using combination therapy with AZA and checkpoint inhibitors have shown some clinical efficacy, and it has been postulated that the antiviral response induced by AZA can sensitize various cancers (7, 17). Given the common functional impact among the different subtypes of AML cell lines as well as cervical (12, 13) and colorectal (14) cancer, combination therapy with AZA and checkpoint inhibitor is a promising strategy for cancer types of hematopoietic origin.

Azacitidine and other DNA methyltransferase inhibitors (DNMTi’s) have been approved for treatment of MDS and AML for the past decade; however, unmet medical need remains for these patients. Here, we used a multi-omics approach to detail the impact of AZA on AML at the individual gene level as well as the functional pathway level. The heterogeneous response of AZA treatment reflects the heterogeneity of cell types, implicating that a sub-type specific therapeutic strategy would be more suitable than a general antibody-based therapy against all AML. Candidate targets for therapeutic targeting in combination with AZA include TRPM4, ADGRE3 and CR1, and validation of these targets in primary patient samples is warranted.

## Methods

### Cell culture and azacitidine treatment

AML193, HL60, KG1a, and HNT34 cells were cultured in RPMI SILAC media (Thermo Fisher Scientific; Waltham, MA) containing L-[^12^C_6_,^14^N_2_]lysine and L-[^12^C_6_,^14^N_4_]arginine (light label) and treated with vehicle (DMSO) or 0.5 μM AZA daily for 3 days. Cells were cultured for another 4 days in drug-free media before frozen cell pellets were harvested for DNA methylation (Infinium MethylationEPIC BeadChip, Illumina) and gene expression (GeneChip^TM^ Human Genome U133 Plus 2.0 Array, Thermo Fisher Scientific) analyses.

### DNA methylation and gene expression data analysis

For DNA methylation data, the standard beta values from Illumina’s BeadStudio were used as our initial input and further corrected for different probe designs using Beta Mixture Quantile dilation (BMIQ) method (35). Probes with known SNPs and too many missing values were also removed from subsequent analyses. The final input data matrix contains 833622 probes. For differential methylation analyses, beta values were transformed to M value (a logit transformation of beta values) and limma package (36) was applied to assess the significance.

For gene expression data, gene expression values were calculated based on the Robust Multi-array Average (RMA) algorithm (37). For each gene, a single probe set was selected based on jetset algorithm (38) resulting the final input data matrix of 20517 genes. Functional enrichment of baseline gene expression profiles was assessed using GSVA package from Bioconductor (39). The package limma was used to assess differential gene expression (36) and gene set enrichment analysis was carried out using fast pre-ranked gene set enrichment analysis (fgsea) package from Bioconductor (40).

Hierarchical clustering based on Euclidean distance and Ward’s minimum variance method was used to evaluate the similarity/difference between samples based on their global DNA methylation or gene expression profiles. The integrated analyses combining DNA methylation and gene expression data were carried out using SMITE package in Bioconductor (34).

All data analyses were carried out using R: A language and environment for statistical computing (41).

### SILAC surface proteomics sample preparation

For stable isotope labeling with amino acids in cell culture (SILAC) experiments, each cell line was cultured in RPMI SILAC media (Thermo Fisher Scientific) containing L-[^13^C_6_,^15^N_2_]lysine and L-[^13^C_6_,^15^N_4_] arginine (heavy label) (Thermo Fisher Scientific) or L-[^12^C_6_,^14^N_2_]lysine and L-[^12^C_6_,^14^N_4_]arginine (light label) for 5 passages to ensure full incorporation of the isotope labeling on cells. In the forward SILAC experiment, heavy labeled cells were treated with AZA while the light labeled cells were treated with DMSO control. In a parallel reverse SILAC experiment, light labeled cells were treated with AZA while the heavy labeled cells were treated with DMSO control. On day 7 of the aforementioned drug treatment, 40-60 × 10^6^ cells from AZA or DMSO treated cells were mixed at a 1:1 cell count ratio for cell surface capture enrichment with several optimization (27). Briefly, live cells were treated with a sodium periodate buffer (2 mM NaPO_4_, PBS pH 6.5) at 4°C for 20 mins to oxidize terminal sialic acids of glycoproteins. Aldehydes generated by periodate oxidation were then reacted with biocytin hydrazide in a labeling buffer (1 mM biocytin hydrazide (biotium), 10 mM analine (Sigma), PBS pH 6.5) at 4°C for 90 mins. Cells were then washed four times in PBS pH 6.5 to remove excess biocytin-hydrazide and flash frozen.

Frozen cell pellets were lysed using RIPA buffer (VWR) with protease inhibitor cocktail (Sigma-Aldrich; St. Louis, MO) at 4°C for 30 mins. Cell lysate was then sonicated, clarified, and incubated with 500μL of neutravidin agarose slurry (Thermo Fisher Scientific) at 4°C for 30 mins. The neutravidin beads were then extensively washed with RIPA buffer, high salt buffer (1M NaCl, PBS pH 7.5), and urea buffer (2M urea, 50mM ammonium bicarbonate) to remove non-specific proteins. Samples were then reduced on-bead with 5mM TCEP at 55°C for 30 mins and alkylated with 10mM iodoacetiamide at room temperature for 30 mins. To release bound proteins, we first performed an on-bead digestion using 20μg trypsin (Promega; Madison, WI) at room temperature overnight. The “tryptic” fraction was then eluted using spin column and the neutravidin beads were extensively washed again with RIPA buffer, high salt buffer (1M NaCl, PBS pH 7.5), and urea buffer (2M urea, 50mM ammonium bicarbonate). To release the remaining trypsin digested N-glycosylated peptides bound to the neutravidin beads, we performed a second on-bead digestion using 2500U PNGase F (New England Biolabs; Ipswich, MA) at 37°C for 3 hrs. Similarly, the “PNGase F” fraction was eluted using a spin column. Both tryptic and PNGase F fractions were then desalted using SOLA HRP SPE column (Thermo Fisher Scientific) using standard protocol, dried, and dissolved in 0.1% formic acid, 2% acetonitrile prior to LC-MS/MS analysis.

### Mass spectrometry analysis

Approximately 1μg of peptide was injected to a pre-packed 0.75mm x 150mm Acclaimed Pepmap C18 reversed phase column (2μm pore size, Thermo Fisher Scientific) attached to a Q Exactive Plus (Thermo Fisher Scientific) mass spectrometer. For “tryptic” fraction, peptides were separated using a linear gradient of 3-35% solvent B (Solvent A: 0.1% formic acid, solvent B: 80% acetonitrile, 0.1% formic acid) over 180 mins at 300μL/min. Similarly, the “PNGase F” fraction was separated using the same gradient over 120 mins. Data were collected in data-dependent mode using a top 20 method with dynamic exclusion of 35 secs and a charge exclusion setting that only sample peptides with a charge of 2, 3, or 4. Full (ms1) scans spectrums were collected as profile data with a resolution of 140,000 (at 200 *m*/*z*), AGC target of 3E6, maximum injection time of 120 ms, and scan range of 400 *-* 1800 *m*/*z*. MS-MS scans were collected as centroid data with a resolution of 17,500 (at 200 *m*/*z*), AGC target of 5E4, maximum injection time of 60 ms with normalized collision energy at 27, and an isolation window of 1.5 *m*/*z* with an isolation offset of 0.5 *m*/*z*.

### Proteomics data processing

Peptide search for each individual dataset was performed using ProteinProspector (v5.13.2) against 20203 human proteins (Swiss-prot database, obtained March 5, 2015) with a false discovery rate (FDR) of <1%. To estimate the efficiency of the surface proteome enrichment method, a list of extracellular proteins was generated by searching for “membrane” but not “mitochondrial” or “nuclear” using uniprot subcellular localization annotations. We found ~60% of peptides identified in the tryptic fraction and ~90% of peptides identified in the PNGase F fraction belonged to the extracellular proteome reflecting a high and expected enrichment ratio.

Quantitative data analysis was performed using Skyline (UWashington) software using the ms1 filtering function. Specifically, spectral libraries from forward and reverse SILAC experiments were analyzed together such that ms1 peaks without an explicit peptide ID would be quantified based on aligned peptide retention time. An isotope dot product of at least 0.8 was used to filter out low quality peptide quantification, and a custom report from skyline was then exported for further processing and analysis using R. To ensure stringent quantification of the surface proteome, several filters were applied to eliminate low confidence protein identifications. In the tryptic fraction, only peptides with five or more well quantified peptides were included. In the PNGase F fraction, only peptides N to D deamidation modification were included. Forward and reverse SILAC datasets were then combined and reported as median log_2_ enrichment values for the AZA treated cells.

## Acknowledgements

This study was funded by the Celgene Corporation. KKL was funded by the CIHR Postdoctoral Fellowship Award. JAW was funded by NIH Grants R35GM122451, P41CA196276 and the Chan Zuckerberg Biohub Investigator award program.

## Competing Interest

This study was funded by the Celgene Corporation. AN, TS, LT, XN, LE, KJM, and JD are employees of Celgene Corporation. JAW and KKL received research funding from Celgene Corporation.

## Supplemental Figures

**Supplemental Figure 1.**
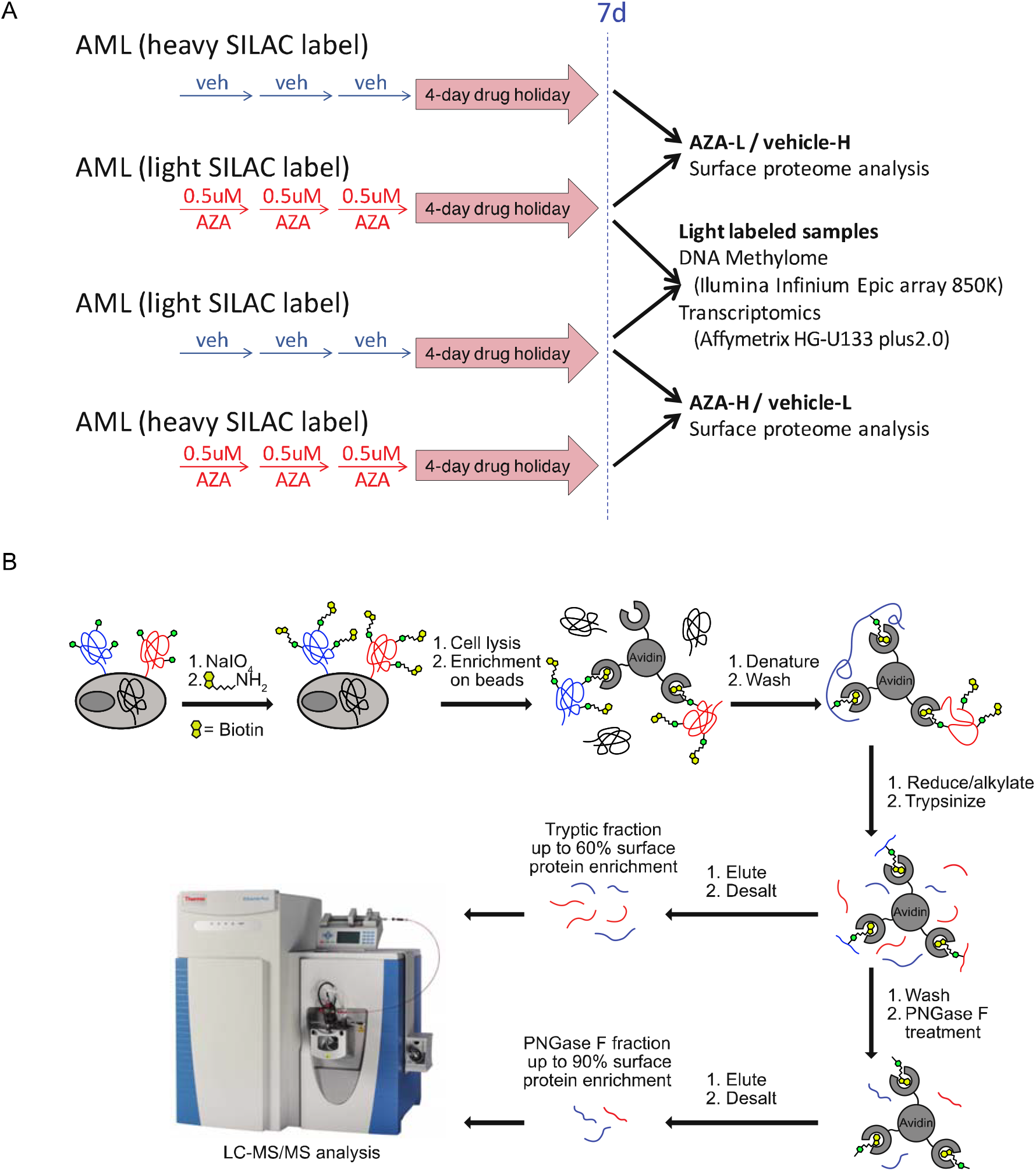
Azacitidine treatment schematic and surface proteome capture methods. (A) The AML cell lines were treated with 0.5μM fresh AZA or vehicle for three days and allowed to recover for four days. The AML AZA-vehicle pairs were grown in either heavy SILAC or light SILAC media. AML cell lines tested in the current study include KG1a, HL60, HNT34, and AML193. (B) Cell surface capture method by biocytin-hydrazide labeling of glycoproteins (see methods for detailed descriptions).

**Supplemental Figure 2.**
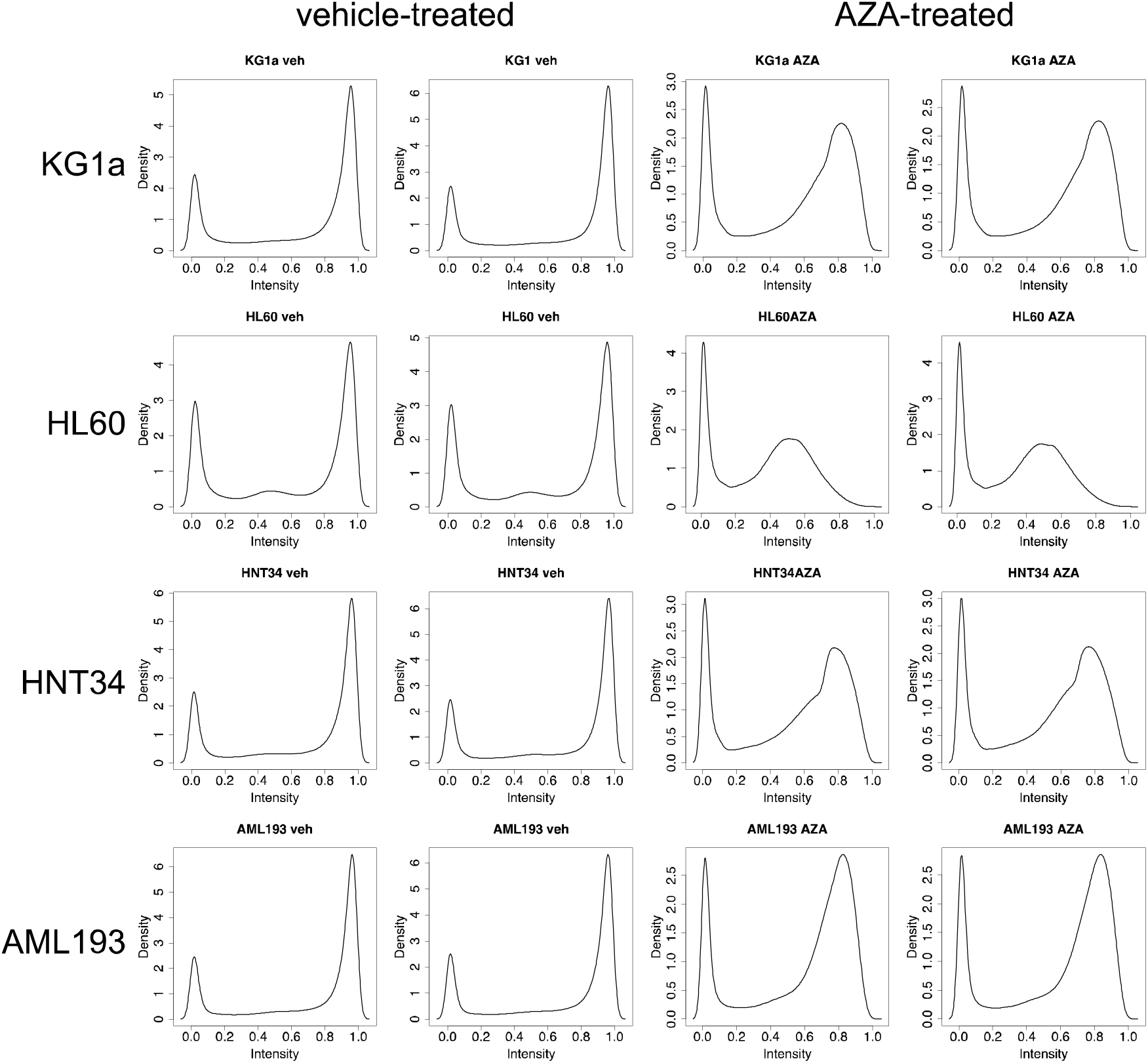
Global de-methylation of AML cell lines when treated by AZA. Density plot of beta values for each cell line sample before and after treatment indicate a general shift of the hypermethylation peak.

**Supplemental Figure 3.**
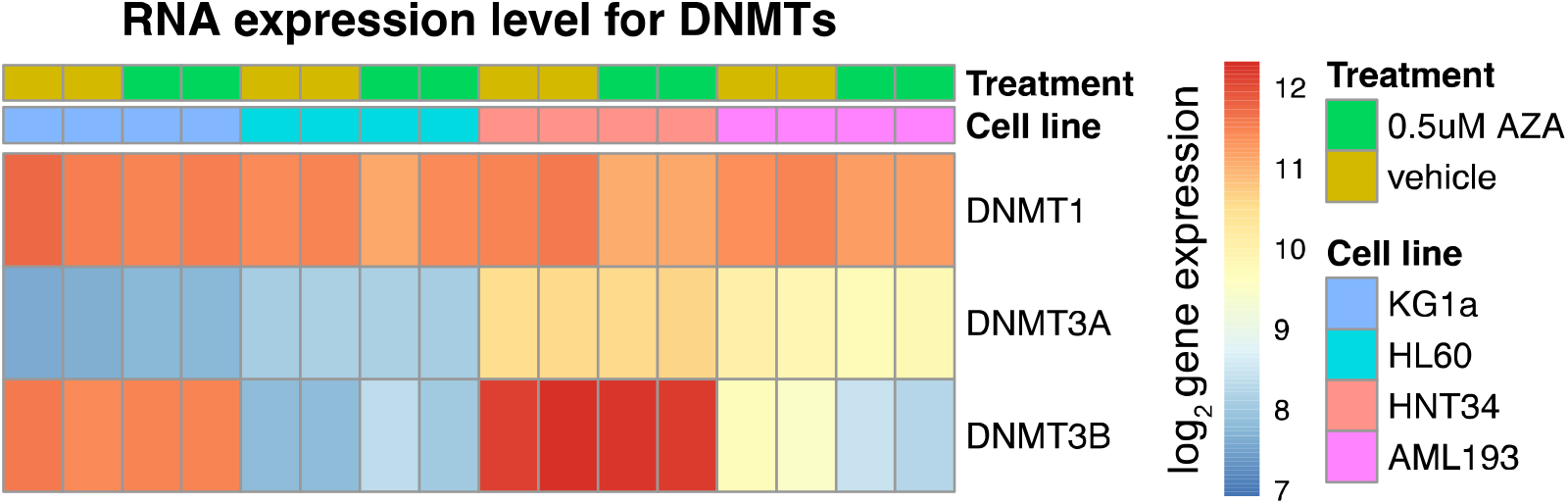
Expression of DNMT transcript in each cell line. Expression of DNMT1 transcript is high in all four cell lines. Expression of both *de novo* methyltransferases DMNT3A and DMNT3B were low in HL60.

**Supplemental Figure 4.**
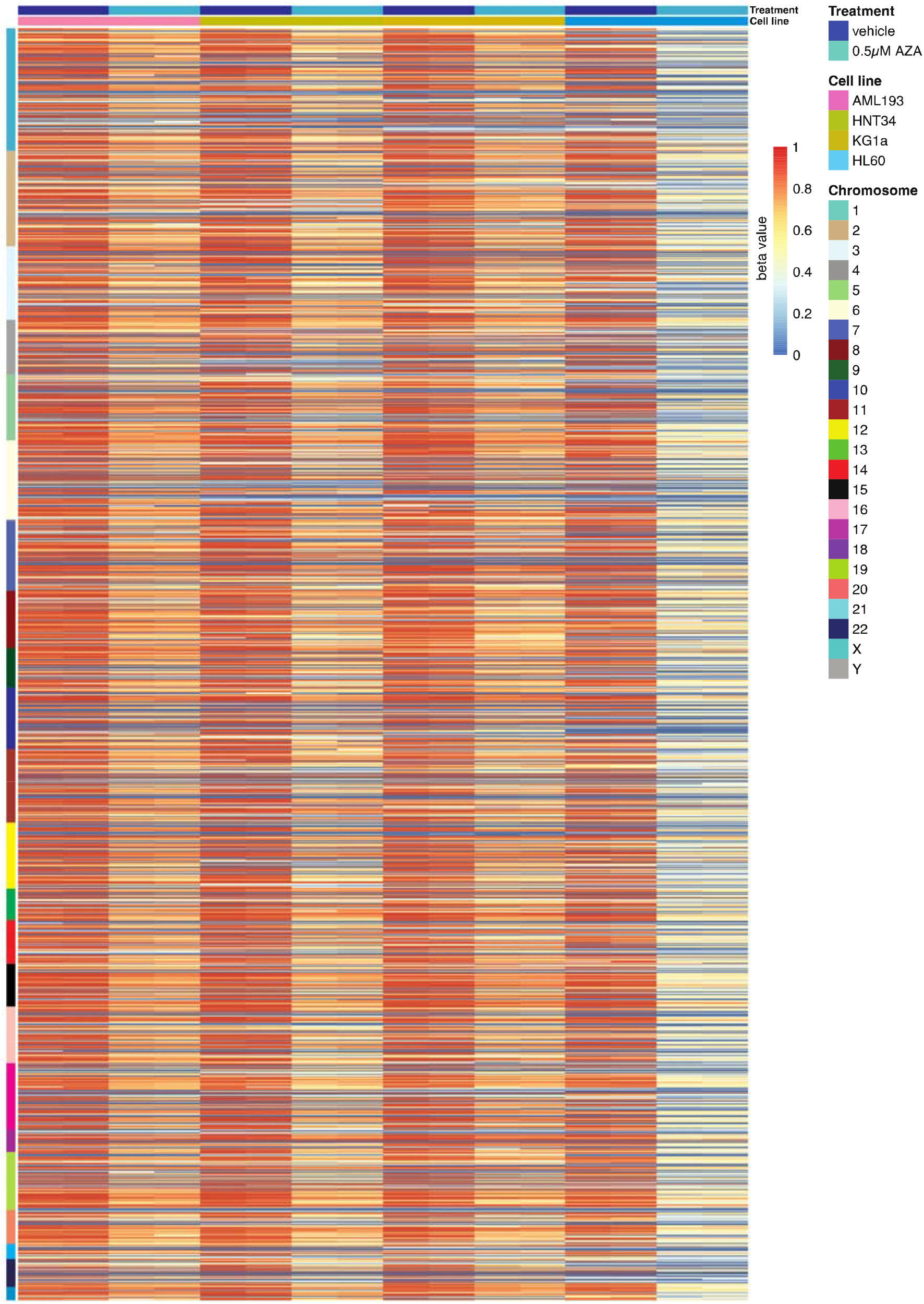
Heatmap of DNA methylation level (beta values) of all ~800,000 CpG arranged by their chromosomal locations in all samples.

**Supplemental Figure 5.**
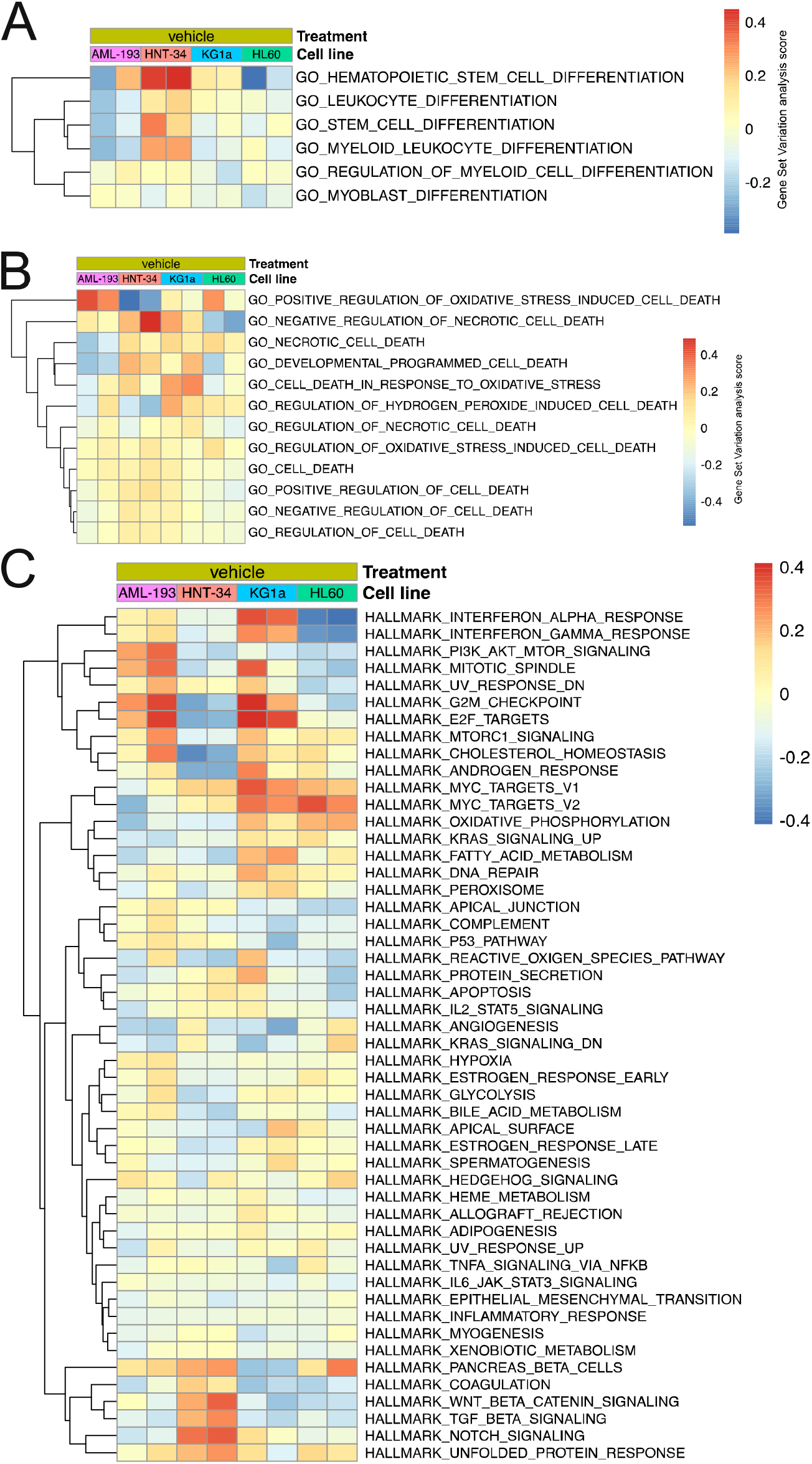
Baseline gene set variation analysis by gene expression data. (A-B) GSVA using Gene Ontology (GO) term analysis in differentiation and cell death pathways indicate differences between the cell lines at baseline. (C) GSVA of the top 50 hallmark gene set indicate different biological states between each cell line.

**Supplemental Figure 6.**
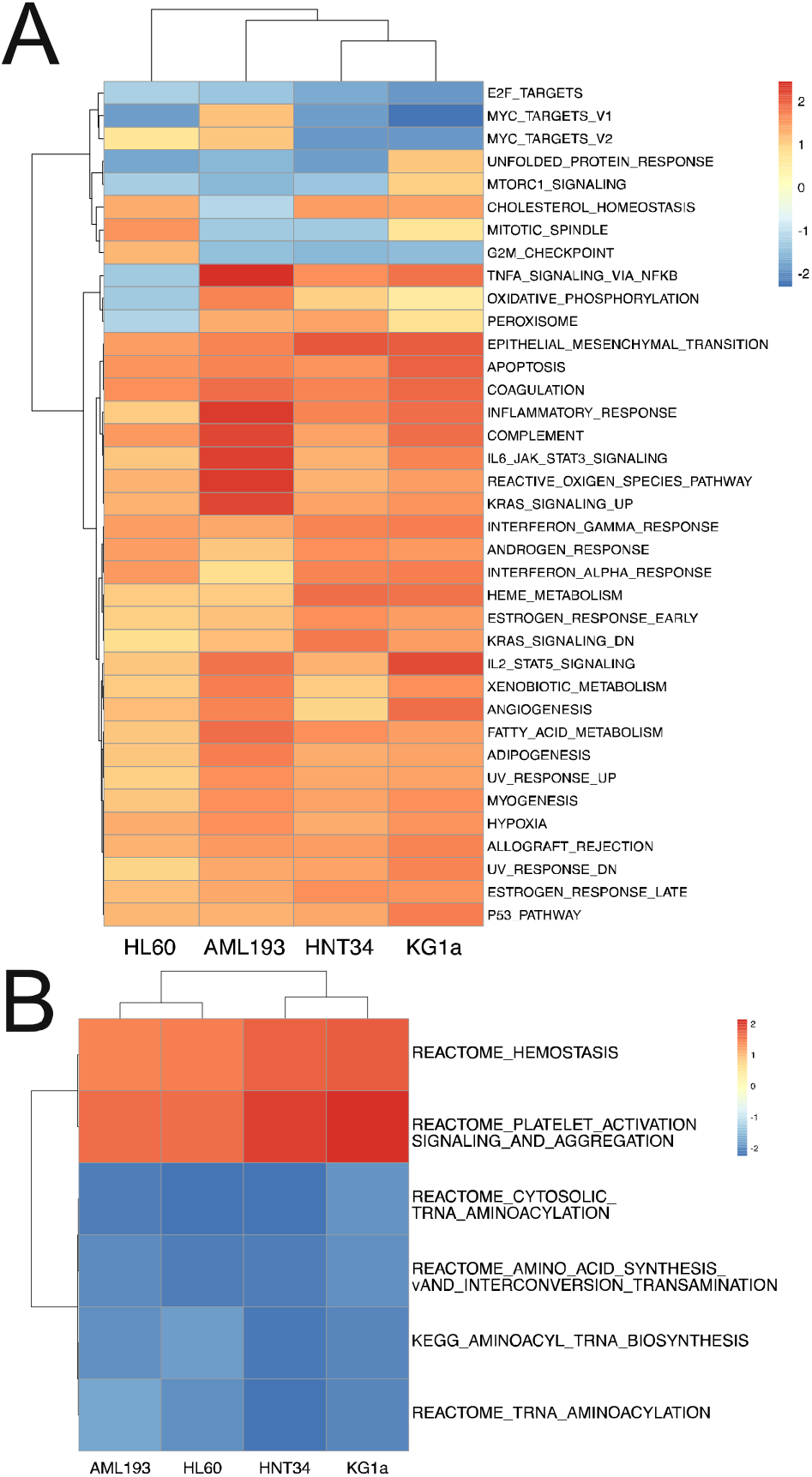
GSEA of gene expression data reveals common biologically processes induced by AZA. (A-B) Gene set enrichment analysis (GSEA) of differentially expressed genes (adjusted p value < 0.05) using hallmark (A), reactome and KEGG gene sets (B).

**Supplemental Figure 7.**
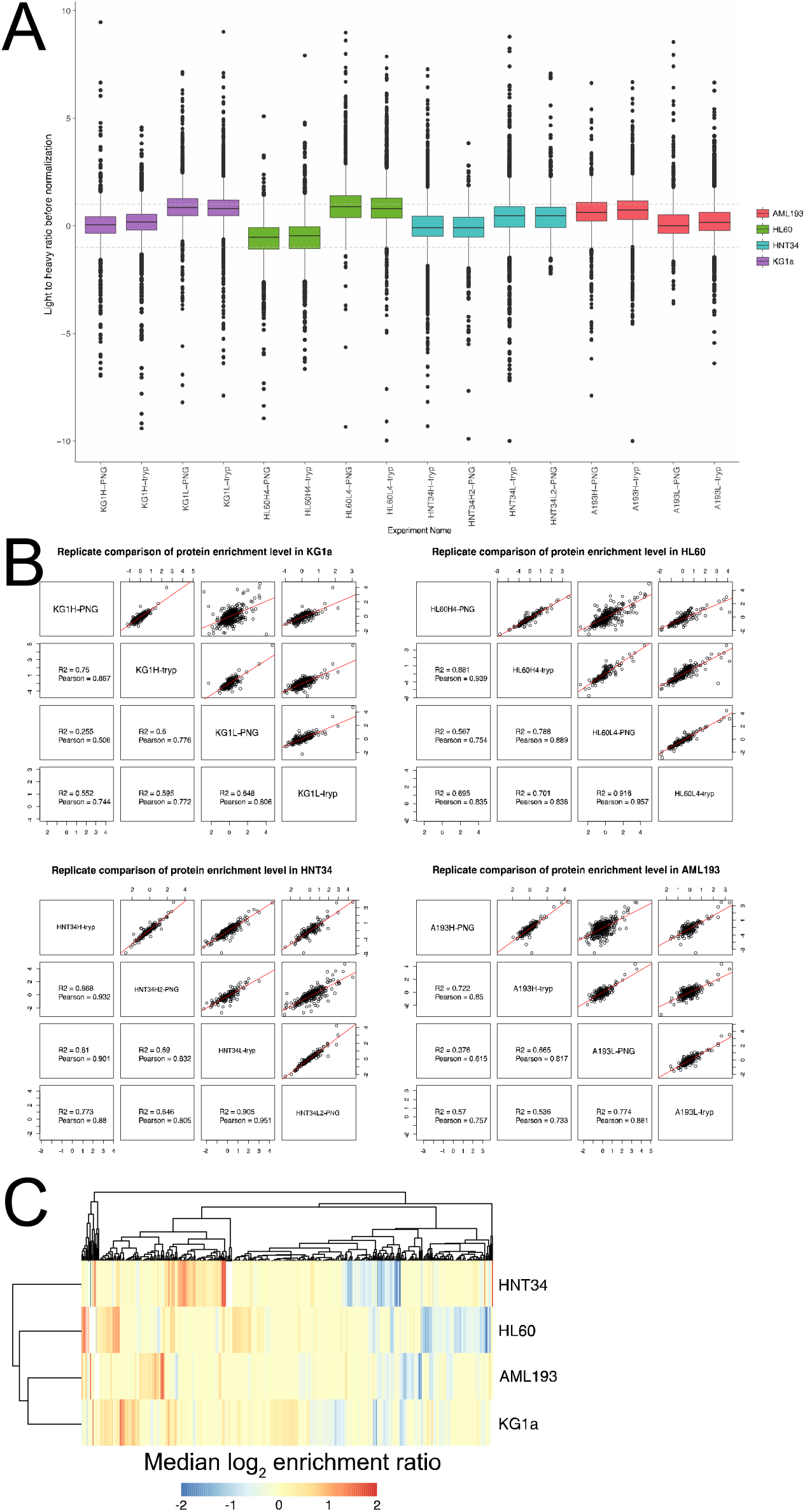
Proteomics analysis statistics. (A) Boxplot of light/heavy SILAC ratio for all datasets. (B) Replicate comparison between each cell line and each fraction. (C) Hierarchal clustering of log_2_ SILAC surface protein enrichment ratio showed a distinct response across the four cell lines when treated with AZA and did not correspond to lineage marker similarity.

**Supplemental Figure 8.**
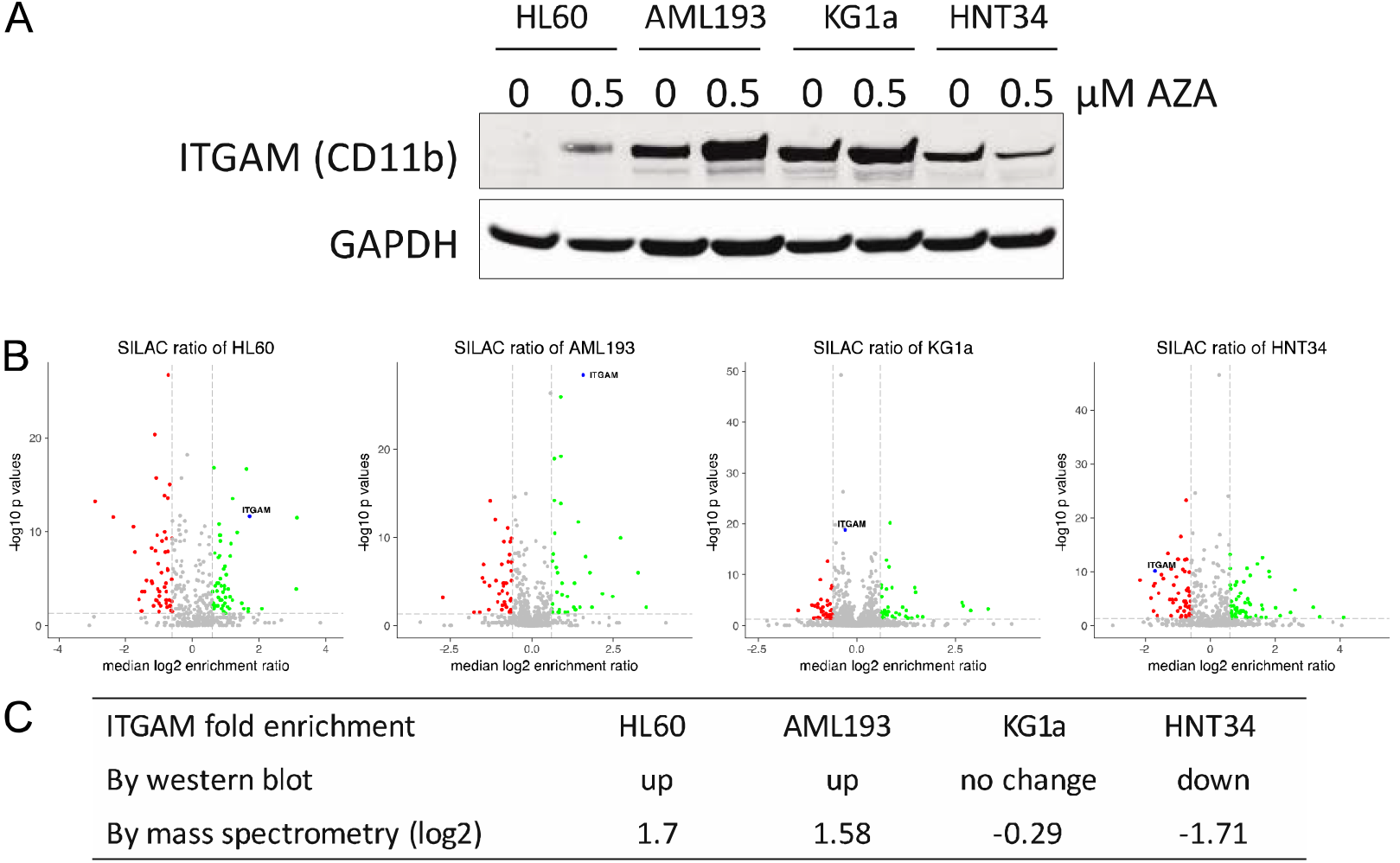
Validation of ITGAM regulation by western blot. (A) Western blot of ITGAM in each cell line with vehicle or AZA treatment indicate different response in each cell line. (B) Volcano plots of surface proteomics data with ITGAM labeled in blue in each cell line. (C) Summary of ITGAM regulation indicating up-regulation in HL60 and AML193 cells, no change in KG1a cells and is down-regulation in HNT34 cells.

**Supplemental Figure 9.**
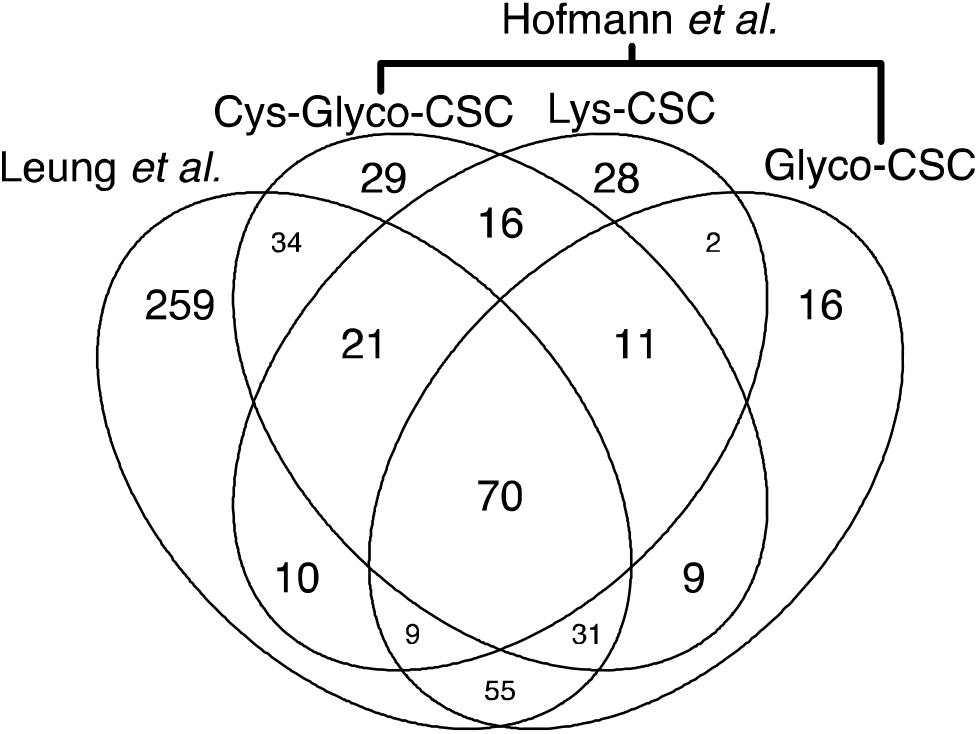
Comparison of surface proteomics data on HL60 cells. Overlap of proteins identified in current study compared to those identified by Hofmann *et al.* (18). A total of 230 proteins were commonly identified in both studies.

**Supplemental Figure 10.**
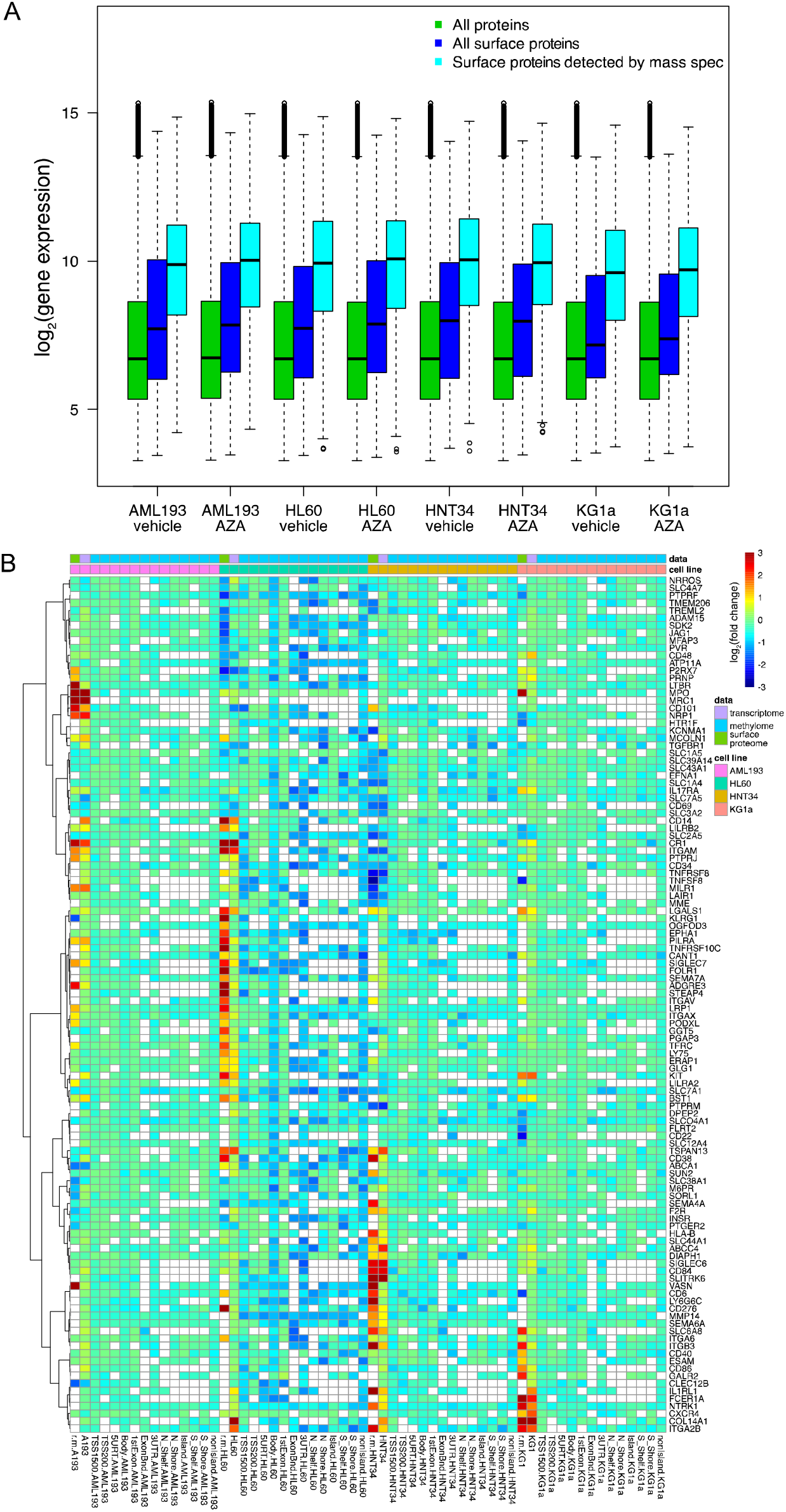
(A) Comparison of log_2_ RNA expression profile of all genes (green), of all genes annotated to be surface proteins (blue), and of genes identified by mass spectrometry experiment (cyan) across all cell lines. Whiskers extends up to 1.5 interquartile range above or below the box. (B) Extension to Figure 4D, showing changes in protein, gene expression, and DNA methylation for all surface protein observed are shown in a heatmap. In addition to TSS1500, methylation changes in other regions such as TSS200, 5’UTR, gene body, 1^st^ exon, exon boundary, 3’UTR, north shelf, north shore, CpG island, south shore, south shelf, and non CpG island are also included.

**Supplemental Figure 11.**
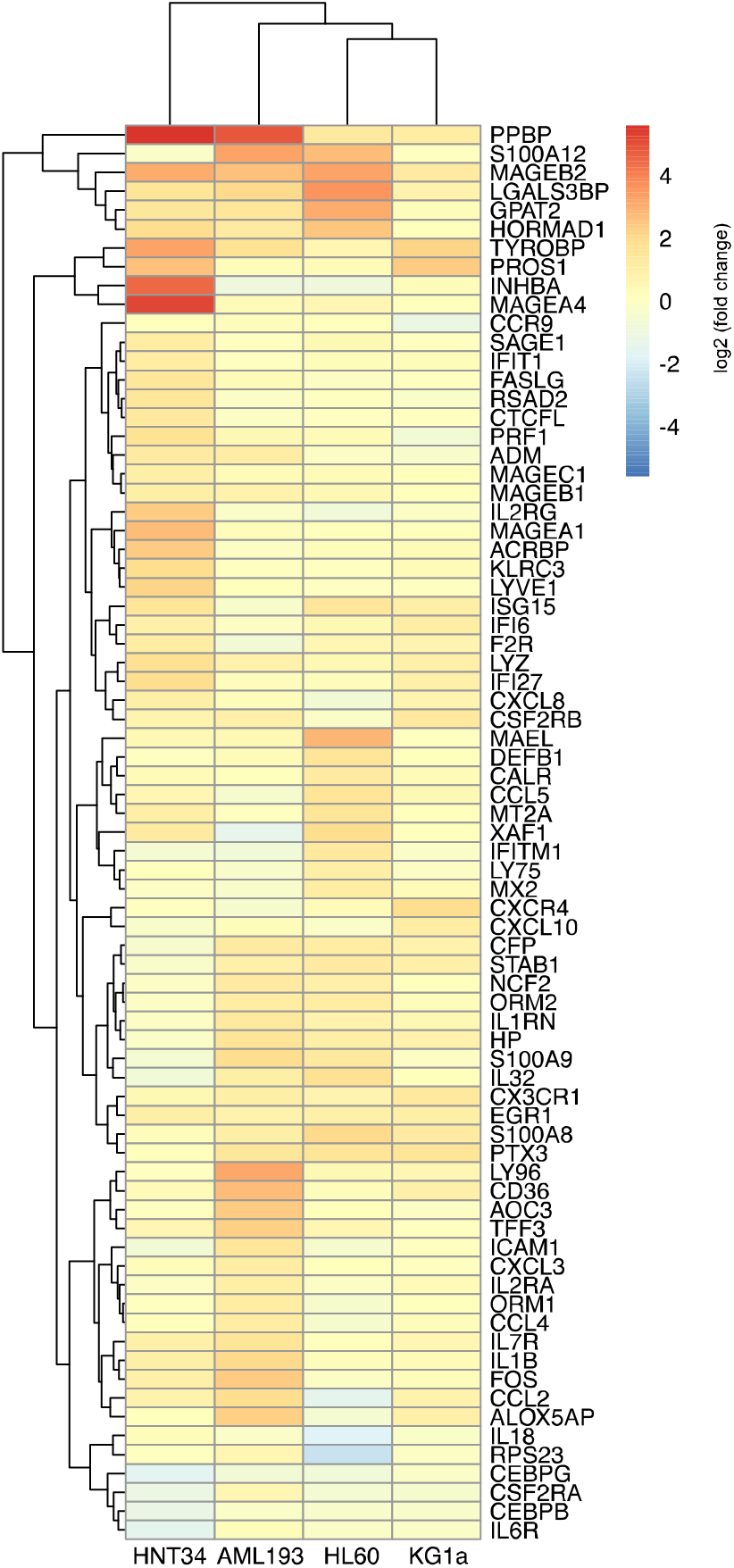
Expression of specific transcripts in the AIM gene set shows an overall enrichment of the AIM gene set to various degree.

## Notes

Conflict of Interest: This study was funded by the Celgene Corporation. AN, TS, LT, XN, LE, KJM, and JD are employees of Celgene Corporation. JAW and KKL received research funding from Celgene Corporation.

## References

1. Issa JJ. The myelodysplastic syndrome as a prototypical epigenetic disease. Blood 2013; 121: 3811-7.

2. Döhner H, Estey E, Grimwade D, Amadori S, Appelbaum FR, Büchner T, et al. Diagnosis and management of AML in adults: 2017 ELN recommendations from an international expert panel. Blood 2017; 129: 424-47.

3. Döhner H, Estey EH, Amadori S, Appelbaum FR, Büchner T, Burnett AK, et al. Diagnosis and management of acute myeloid leukemia in adults: recommendations from an international expert panel, on behalf of the European LeukemiaNet. Blood 2010; 115: 453-74.

4. Allis CD, Jenuwein T. The molecular hallmarks of epigenetic control. Nat Rev Genet 2016; 17: 487-500.

5. Esteller M. Cancer epigenomics: DNA methylomes and histone-modification maps. Nat Rev Genet 2007; 8: 286-98.

6. Mund C, Brueckner B, Lyko F. Reactivation of epigenetically silenced genes by DNA methyltransferase inhibitors: basic concepts and clinical applications. Epigenetics 2006; 1: 7-13.

7. Wolff F, Leisch M, Greil R, Risch A, Pleyer L. The double-edged sword of (re)expression of genes by hypomethylating agents: from viral mimicry to exploitation as priming agents for targeted immune checkpoint modulation. Cell Commun Signal 2017; 15: 13.

8. Dombret H, Seymour JF, Butrym A, Wierzbowska A, Selleslag D, Jang JH, et al. International phase 3 study of azacitidine vs conventional care regimens in older patients with newly diagnosed AML with >30% blasts. Blood 2015; 126: 291-9.

9. Silverman LR, Demakos EP, Peterson BL, Kornblith AB, Holland JC, Odchimar-Reissig R, et al. Randomized controlled trial of azacitidine in patients with the myelodysplastic syndrome: a study of the cancer and leukemia group B. J Clin Oncol 2002; 20: 2429-40.

10. Stresemann C, Lyko F. Modes of action of the DNA methyltransferase inhibitors azacytidine and decitabine. Int J Cancer 2008; 123: 8-13.

11. Tsai H, Li H, Van Neste L, Cai Y, Robert C, Rassool FV, et al. Transient low doses of DNA-demethylating agents exert durable antitumor effects on hematological and epithelial tumor cells. Cancer Cell 2012; 21: 430-46.

12. Li H, Chiappinelli KB, Guzzetta AA, Easwaran H, Yen RC, Vatapalli R, et al. Immune regulation by low doses of the DNA methyltransferase inhibitor 5-azacitidine in common human epithelial cancers. Oncotarget 2014; 5: 587-98.

13. Chiappinelli KB, Strissel PL, Desrichard A, Li H, Henke C, Akman B, et al. Inhibiting DNA Methylation Causes an Interferon Response in Cancer via dsRNA Including Endogenous Retroviruses. Cell 2015; 162: 974-86.

14. Roulois D, Loo Yau H, Singhania R, Wang Y, Danesh A, Shen SY, et al. DNA-Demethylating Agents Target Colorectal Cancer Cells by Inducing Viral Mimicry by Endogenous Transcripts. Cell 2015; 162: 961-73.

15. Martinko Alexander J, Truillet Charles, Julien Olivier, Diaz Juan E, Horlbeck Max A, Whiteley Gordon, et al. Targeting RAS-driven human cancer cells with antibodies to upregulated and essential cell-surface proteins. eLife 2018; 7.

16. Hoseini SS, Cheung NK. Acute myeloid leukemia targets for bispecific antibodies. Blood Cancer J 2017; 7: e522.

17. Daver N, Boddu P, Garcia-Manero G, Yadav SS, Sharma P, Allison J, et al. Hypomethylating agents in combination with immune checkpoint inhibitors in acute myeloid leukemia and myelodysplastic syndromes. Leukemia 2018; 32: 1094-105.

18. Hofmann A, Gerrits B, Schmidt A, Bock T, Bausch-Fluck D, Aebersold R, et al. Proteomic cell surface phenotyping of differentiating acute myeloid leukemia cells. Blood 2010; 116: 26.

19. Perna F, Berman SH, Soni RK, Mansilla-Soto J, Eyquem J, Hamieh M, et al. Integrating Proteomics and Transcriptomics for Systematic Combinatorial Chimeric Antigen Receptor Therapy of AML. Cancer Cell 2017; 32: 519.e5.

20. Furley AJ, Reeves BR, Mizutani S, Altass LJ, Watt SM, Jacob MC, et al. Divergent molecular phenotypes of KG1 and KG1a myeloid cell lines. Blood 1986; 68: 1101-7.

21. Koeffler HP, Billing R, Lusis AJ, Sparkes R, Golde DW. An undifferentiated variant derived from the human acute myelogenous leukemia cell line (KG-1). Blood 1980; 56: 265-73.

22. Gallagher R, Collins S, Trujillo J, McCredie K, Ahearn M, Tsai S, et al. Characterization of the continuous, differentiating myeloid cell line (HL-60) from a patient with acute promyelocytic leukemia. Blood 1979; 54: 713-33.

23. Yamamoto K, Hamaguchi H, Nagata K, Kobayashi M, Tanimoto F, Taniwaki M. Establishment of a novel human acute myeloblastic leukemia cell line (YNH-1) with t(16;21), t(1;16) and 12q13 translocations. Leukemia 1997; 11: 599-608.

24. Lange B, Valtieri M, Santoli D, Caracciolo D, Mavilio F, Gemperlein I, et al. Growth factor requirements of childhood acute leukemia: establishment of GM-CSF-dependent cell lines. Blood 1987; 70: 192-9.

25. Zhang W, Fei J, Yu S, Shen J, Zhu X, Sadhukhan A, et al. LINC01088 inhibits tumorigenesis of ovarian epithelial cells by targeting miR-24-1-5p. Sci Rep 2018; 8: 2876.

26. Ong S, Blagoev B, Kratchmarova I, Kristensen DB, Steen H, Pandey A, et al. Stable isotope labeling by amino acids in cell culture, SILAC, as a simple and accurate approach to expression proteomics. Mol Cell Proteomics 2002; 1: 376-86.

27. Wollscheid B, Bausch-Fluck D, Henderson C, O’Brien R, Bibel M, Schiess R, et al. Mass-spectrometric identification and relative quantification of N-linked cell surface glycoproteins. Nat Biotechnol 2009; 27: 378-86.

28. Valtieri M, Boccoli G, Testa U, Barletta C, Peschle C. Two-step differentiation of AML-193 leukemic line: terminal maturation is induced by positive interaction of retinoic acid with granulocyte colony-stimulating factor (CSF) and vitamin D3 with monocyte CSF. Blood 1991; 77: 1804-12.

29. Lundberg E, Fagerberg L, Klevebring D, Matic I, Geiger T, Cox J, et al. Defining the transcriptome and proteome in three functionally different human cell lines. Mol Syst Biol 2010; 6: 450.

30. Björn Schwanhäusser, Dorothea Busse, Na Li, Gunnar Dittmar, Johannes Schuchhardt, Jana Wolf, et al. Global quantification of mammalian gene expression control. Nature 2011; 473: 337-42.

31. Wiita AP, Ziv E, Wiita PJ, Urisman A, Julien O, Burlingame AL, et al. Global cellular response to chemotherapy-induced apoptosis. Elife 2013; 2: e01236.

32. Klco JM, Spencer DH, Lamprecht TL, Sarkaria SM, Wylie T, Magrini V, et al. Genomic impact of transient low-dose decitabine treatment on primary AML cells. Blood 2013; 121: 1633-43.

33. Yin Y, Morgunova E, Jolma A, Kaasinen E, Sahu B, Khund-Sayeed S, et al. Impact of cytosine methylation on DNA binding specificities of human transcription factors. Science 2017; 356.

34. Wijetunga NA, Johnston AD, Maekawa R, Delahaye F, Ulahannan N, Kim K, et al. SMITE: an R/Bioconductor package that identifies network modules by integrating genomic and epigenomic information. BMC Bioinformatics 2017; 18: 41.

35. Teschendorff AE, Marabita F, Lechner M, Bartlett T, Tegner J, Gomez-Cabrero D, et al. A beta-mixture quantile normalization method for correcting probe design bias in Illumina Infinium 450 k DNA methylation data. Bioinformatics 2013; 29: 189-96.

36. Smyth GK. Linear models and empirical bayes methods for assessing differential expression in microarray experiments. Stat Appl Genet Mol Biol 2004; 3: Article 3.

37. Irizarry RA, Hobbs B, Collin F, Beazer-Barclay YD, Antonellis KJ, Scherf U, et al. Exploration, normalization, and summaries of high density oligonucleotide array probe level data. Biostatistics 2003; 4: 249-64.

38. Li Q, Birkbak NJ, Gyorffy B, Szallasi Z, Eklund AC. Jetset: selecting the optimal microarray probe set to represent a gene. BMC Bioinformatics 2011; 12: 474.

39. Hänzelmann S, Castelo R, Guinney J. GSVA: gene set variation analysis for microarray and RNA-seq data. BMC Bioinformatics 2013; 14: 7.

40. Alexey A. Sergushichev. An algorithm for fast preranked gene set enrichment analysis using cumulative statistic calculation.

41. R Core Team. R: A language and environment for statistical computing. R Foundation for Statistical Computing, Vienna, Austria. 2018.

